# Early phyllosphere microbial associations impact plant reproductive success

**DOI:** 10.1101/2022.06.30.498294

**Authors:** Elijah C. Mehlferber, Kent F. McCue, Reena Debray, Griffin Kaulbach, Jon E. Ferrel, Rajnish Khanna, Britt Koskella

## Abstract

- The above-ground plant microbiome (the phyllosphere) is increasingly recognized as an important component of plant health. We hypothesized that phyllosphere interactions may be disrupted in a greenhouse setting, where microbial dispersal is limited, and that adding a microbial amendment might yield important benefits to the host plant.
- Using a newly developed synthetic phyllosphere microbiome for Tomato, we tested this hypothesis across multiple trials by manipulating microbial colonization of leaves and measuring subsequent plant growth and reproductive success, comparing results from plants grown in both greenhouse and field settings.
- We confirmed that greenhouse-grown plants have a depauperate phyllosphere microbiome and that the addition of the synthetic microbial community was responsible for a clear and repeatable increase in fruit production in this setting. We further show that this effect is synergistic with the addition of micronutrient-based soil amendments, with important implications for agriculture.
- These results suggest that greenhouse environments have poor phyllosphere microbiome establishment, with negative impacts on the plant. The results also implicate the phyllosphere microbiome as a key component of plant fitness, emphasizing that these communities have a clear role to play in the ecology and evolution of plant communities.

## Introduction

Microbial associations have been shown to be critical in the development and functioning of plant and animal host organisms (Shin *et al*., 2011; Wagner *et al*., 2014). For plants, there exists a wealth of data on how the root and soil-associated microbial communities can shape plant growth, competition with neighbors, disease resistance, and nutrient uptake (Berendsen *et al*., 2012). In contrast to the well-defined role of belowground plant-associated communities, less is known about the importance of bacteria inhabiting the above-ground portion of the plant, the phyllosphere. Thus far, the investigation of phyllosphere microbiomes has generally been limited to their role in protection from disease, such as in cases of pear fire blight (Mercier & Lindow, 2001), tobacco wildfire disease (Qin *et al*., 2019), or tomato bacterial speck (Innerebner *et al*.,2011; Morella *et al*., 2019). Although some evidence suggests that these microbial communities can have key functions beyond disease resistance (reviewed in Stone *et al*., 2018), for example through nitrogen fixation (Fürnkranz *et al*., 2008) or the production of growth-regulating signals (Madhaiyan *et al*., 2006), there remains limited direct evidence for their role in plant growth or yield.

The phyllosphere is inhabited by a relatively diverse consortia of bacteria, with densities ranging from 10^6 to 10^7 cells per square centimeter (Lindow & Brandl, 2003). These epiphytic bacteria are subject to a hostile environment, often encountering high levels of UV radiation, temperature fluctuations, and desiccation (Jacobs *et al*., 2005; Beattie, 2011). The majority of phyllosphere-inhabiting bacteria are believed to arrive via aerial transmission, including wind and rain (Vorholt, 2012; Ottesen *et al*., 2016), and much of this transmission is likely to originate from neighboring plants (Šantl-Temkiv *et al*., 2018; Meyer *et al*., 2022). In contrast, plants that are grown in greenhouses are relatively isolated from microbial dispersal through wind, rain, and insects or from neighboring plants. In line with this, greenhouse-grown plants have been shown to develop communities distinct from those developing in outdoor environments (Maignien *et al*., 2014). Since greenhouse plants are typically grown in commercial soil mixes, it is also unlikely that they have the full breadth of bacteria available for recruitment from the soil reservoir (Knief *et al*., 2010). Given this, greenhouse-grown plants are likely depauperate in their microbial associations, providing a unique opportunity to study the importance of the phyllosphere microbiota in a more complex environment, as opposed to a highly constrained gnotobiotic system.

One promising avenue for investigating the causative effects of plant-microbiota interactions in plant health is through the use of synthetic bacterial communities. Ideally, these synthetic communities represent the phylogenetic diversity of natural phyllosphere communities, but at a tractable level of complexity, allowing for repeatable experimentation. This approach has been used to investigate a wide variety of plant-microbial interactions (Bodenhausen *et al*., 2014; Bai *et al*., 2015; Hu *et al*., 2016; Castrillo *et al*., 2017; Berg & Koskella, 2018), but these synthetic communities also hold great potential as microbial ‘probiotics’ or biostimulants. Such microbial amendments would be especially useful in environments where microbial diversity is otherwise reduced and/or where host-microbiome associations have been disrupted by, for example, pathogen establishment or antimicrobial treatments. We examine this question using a defined set of naturally occurring bacteria to establish a synthetic community (herein referred to as ‘PhylloStart’) that we developed to mimic the composition of microbial communities associated with field-grown tomato plants.

Our primary objective was to determine the importance of phyllosphere-associated microbiota in agriculturally relevant plant traits, under realistic conditions, and so we focused on early microbiome establishment in a greenhouse setting, where we show that initial microbiome recruitment is highly limited. To investigate potential interactions between microbial associations and nutrient status we included a popular commercially available micronutrient supplement (Azomite) at various concentrations. Over a series of trials we demonstrate both long-term establishment of the synthetic microbiome on plants and a highly repeatable and significant increase in plant growth and yield compared to control plants, with an additive effect from the micronutrient supplement. In the case of field-grown plants that were treated in the same way (i.e., sprayed pre-transplantation with PhylloStart), however, we did not see a significant effect of early life phyllosphere amendment. This differential effect of the phyllosphere on plant fitness across the greenhouse and field is likely the result of reduced microbiome colonization in low diversity/dispersal environment such as the greenhouse. Moreover, higher levels of competition and environmental pressures under field conditions could reduce the efficacy of early life application of probiotic treatments, potentially requiring different application protocols. Overall, our results demonstrate that, typically, phyllosphere microbial communities establish poorly under common greenhouse growth conditions and highlight the underappreciated role of the above-ground microbiome in shaping plant fitness.

## Materials and Methods

### Plant generation

Seeds of tomato *(Solanum lycopersicum)* variety ‘Moneymaker’ were surface sterilized by gently shaking in a solution of sodium hypochlorite and Tween 20 for 20 min, followed by two rinses with filter-sterilized H_2_O. Seeds were placed into pots containing Sunshine mix number 1 (Sun Gro) soil and germinated in the greenhouse. When the seedlings were 3 to 4 inches tall, they were transplanted into larger pots, and randomly distributed across the greenhouse, where they were grown for the remainder of their development, which was 20 weeks in the first trial, 19 weeks in the second trial and 24 weeks in the third trial. Plants were grown in the greenhouse under controlled conditions with supplemented lights to maintain long days and fans to control high-temperature fluctuations. Liquid nutrient supplementation consisting of Peters Professional 20/20/20 water-soluble fertilizer was applied (1:64□ppm) once per week, as well as a disease suppression program consisting of Floramite and Decathlon at a rate of 1/4 tsp per gallon of water, mixed/agitated, was applied through a controlled sprayer at the rate of 1 to 2□gal per 100 plants. Plants in the field trial were started in the greenhouse and received the same treatment as the plants in the third trial until a time of 7 weeks, when they were moved outside to harden, and subsequently transplanted into the Oxford Tract field at UC Berkeley at 8 weeks. After transplanting into the field plants were watered upon establishment as needed, and then once a week for approximately 6 hours thereafter, and 20-20-20 water-soluble fertilizer was applied during watering at a rate of .93#N/ac.

### Bacterial strains

PhylloStart was designed to mimic the composition of a field grown tomato phyllosphere community, but at reduced complexity to remain tractable. The community was initially designed based on communities sequenced from tomato plants in the student organic garden at UC Davis (Supplemental Table 1), and isolates were collected to be representative at the family level. These isolates were collected directly from the UC Davis Student Farm, and from the endpoint of a greenhouse selection experiment (Morella *et al*., 2020) by plating initially on KB and LB agar plates, followed by MacConkey, and 1% Tryptic Soy agar plates to isolate more fastidious species. The isolates that were selected for inclusion comprise 97.8% of the bacteria that were found at a relative abundance greater than 1% in this dataset, representing the families Enterobacteriaceae, Oxalobacteraceae, Pseudomonadaceae, Bacillaceae, Microbacteriaceae, with the addition of a member of the family Brevibacteriaceae that was identified at a high prevalence on our plated field samples during collection. In total, 16 unique species were selected to comprise PhylloStart (out of the 93 screened isolates), with several members representing species level variation within the selected families. Information on the identity of the PhylloStart synthetic community is available in Supplemental Table 2. Strain identification was performed by sequencing the genomes of each bacteria and matching the sequences for the 16s rRNA using BLAST to publicly available databases on NCBI.

### PhylloStart and Azomite application

For preparation of consortia, all strains were grown individually for three days at 28°C on a media shaker in liquid culture in KB (King’s Broth) nutrient broth (King *et al*., 1954). Cultures were spun down for 10min at 2500g in a centrifuge and the KB supernatant was replaced with fresh KB. The optical density at 600nm (OD600) of each sample was read and the samples were mixed together at densities equal to 0.2. This suspension was frozen in 50/50 KB/Glycerol at −80°C until inoculation. On the day of inoculation, the community was thawed and resuspended in sterile 10Mm MgCl buffer with 0.01% SilWet surfactant. For the second greenhouse trial and the field trial we included two concentrations of the PhylloStart, one at OD600=0.02 and another diluted 100-fold at OD600=0.0002. Plants were inoculated by spraying either this suspension, or for controls, the same but without the addition of PhylloStart, onto both sides of all leaves until runoff. Plant inoculation time varied among experiments but was generally performed for three consecutive weeks starting either two- or three-weeks post seedling emergence (See Fig. **1** for exact timing across experiments).

**Fig 1.**
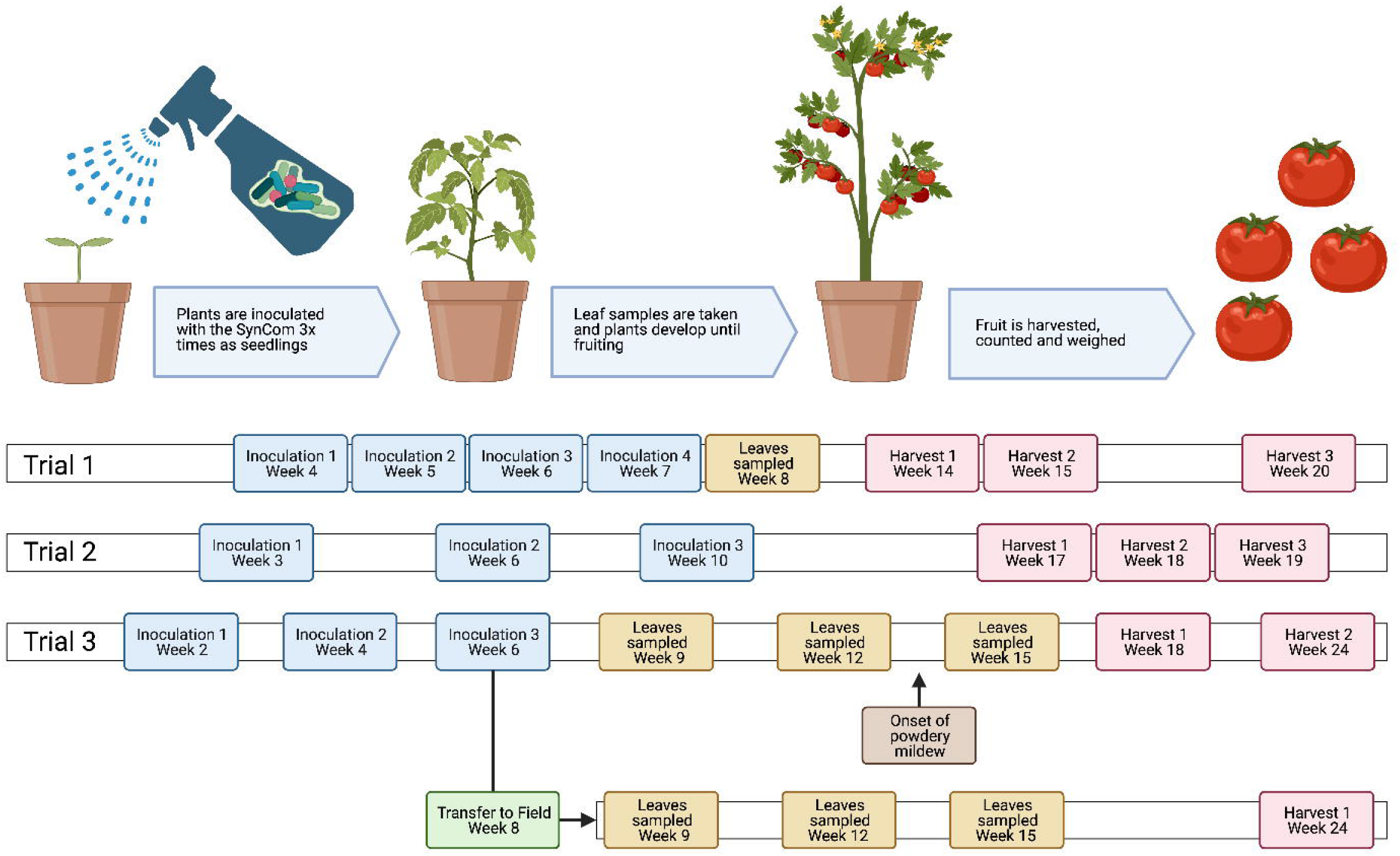
Experimental design for the three greenhouse experiments. Text in Blue indicates the times when the plants were inoculated with the PhylloStart bacteria. Text in Yellow indicates times when leaves were sampled for sequencing. Text in Red indicates timing of tomato harvests. Note the inclusion of the approximate onset of powdery mildew in Trial 3.

The micronutrient supplementation differed between trials, with different preparations and quantities of Azomite®(Azomite Mineral Products, Inc., UT), a soil additive and fertilizer derived from volcanic ash that has previously been shown to increase the growth and yield of tomato plants (Azad *et al*., 2016; Mehlferber *et al*., 2022). There were three trials performed (Fig. **1**). In the first greenhouse trial we used three Azomite treatments, first, with the plants either amended with 5% wt/wt Azomite Granular grade during sowing and transplanting (n = 10), second, with 1g of Azomite Ultrafine grade applied after transplanting to the soil surface at the base of the plant at 7, 9, and 12□weeks after sowing (n = 10), and finally, with both Azomite Granular and Azomite Ultrafine applied as described (n = 10). Trial 1 included a control treatment with no Azomite or PhylloStart (n = 10), as well as a PhylloStart only treatment and a treatment with both PhylloStart and Azomite Granular and Ultrafine (n = 10). In the second greenhouse trial we applied the same Azomite Granular and Ultrafine treatment as described, but with the modification of Azomite Ultrafine concentration, using 1, 2 and 3 grams (n = 3 for each). This experiment included plants that were inoculated with these concentrations of Azomite as well as PhylloStart (n = 3 for each), PhylloStart alone (n = 3), and a control treatment that did not receive either Azomite or PhylloStart (n = 3). In the third greenhouse trial, and the field trial, we included treatments with Azomite Granular and Ultrafine, using 1 and 2 grams in the greenhouse, and 1 and 3 grams in the field (n = 4 for each). Trial 3 included plants that were amended with these concentrations of Azomite and were inoculated (as described in Fig. **1**) with either a low, or high dose of PhylloStart (n = 4 for each), a PhylloStart only treatment at both concentrations (n = 6 for each), and a control treatment that did not receive either Azomite or PhylloStart (n = 6). A second field trial was performed at UC Davis to confirm the results from our initial field trial, using the same methods as previously described, with the following treatments; PhylloStart (n = 6), Control (n = 6), PhylloStart with 3 grams of Azomite (n = 6) and Control with 3 grams of Azomite (n = 6).

### Plant measurements and harvest

Plant height was measured from soil surface to the terminal node, recorded weekly during vegetative growth in Days After Sowing as indicated (Supplemental Fig. **1**). Plant width was determined by measuring the combined lengths of the two longest opposing-side branches at the base of each plant, recorded along with plant height. During reproductive growth, total numbers of flowers and fruit attached per plant were counted as indicated (Supplemental Fig. **1** and Figs. **3, 4**). Tomatoes were weighed individually, and as total harvested weight per plant as described (Supplementary Fig. **2, 3, 4**). Tomato number and weight were recorded multiple times per plant from onset of fruit production to plant termination in the greenhouse, but these metrics were measured only once after harvest from each individual plant grown in the field.

### Leaf sample collection

Leaves were sampled in the first and third experiments to assay the composition of the phyllosphere microbial community. In each case, 5 leaves were collected into 50ml conical tubes from random locations across each plant. These leaves were weighed and 40ml of sterile 10mM MgCl2 was added. They were sonicated for 10 minutes, followed by five seconds of vortexing to ensure that the bacteria separated from the leaves. The bacteria were pelleted, and the supernatant was removed. These samples were frozen at −80°C until DNA extraction and sequencing.

### Pathogen protection experiment

Moneymaker tomato seeds were prepared as described above, then germinated onto plates of 1% water agar. After 1 week, seedlings were transferred to individual pots containing autoclaved soil consisting of calcined clay medium (Profile Porous Ceramic Greens Grade, Sierra Pacific Turf Supply). In the fertilizer treatment, 960 mg of organic fertilizer (0-11-0 Seabird Guano, Down to Earth) was added to each pot at the transplant stage. Plants were randomized with respect to treatment and maintained in a growth chamber at a 15 h day:9 h night cycle for the duration of the experiment.

When plants were three weeks old, PhylloStart communities were applied to leaves at a concentration of OD600=0.02 with 0.01% SilWet surfactant. One week after spraying, an overnight culture of *Pseudomonas syringae* pathovar tomato PT23 was diluted in 10 mM MgCl_2_ to a concentration of OD600=0.0002. Three leaves per plant were challenged via blunt-end syringe inoculation. At 24 hours post-infection, 3 hole punches (6-mm diameter) were taken from each inoculated leaf (9 total leaf discs per plant). Leaf discs were homogenized in 1 mL 10 mM MgCl_2_ in a FastPrep-24 5G sample disruption instrument at 4.0 m/s for 40 seconds. *Pseudomonas syringae* population density on leaves was obtained through colony forming unit (CFU) plating.

### DNA extractions, qPCR, 16s rRNA amplification, and sequencing

DNA extraction and sequencing was performed by Microbiome Insights using the following protocols. Bacterial pellets were placed into a MoBio PowerMag Soil DNA Isolation Bead Plate. DNA was extracted following MoBio’s instructions on a KingFisher robot. For qPCR, bacterial-specific (300 nM 27F, 5’ -AGAGTTTGATCCTGGCTCAG-3’) forward primers coupled to (300 nM 519R, 5’ -ATTACCGCGGCTGCTGG-3’) reverse primers were used to amplify bacterial 16S rRNA. 20 μl reactions using iQ SYBR Green Supermix (Bio□Rad), with 10μl Supermix, 0.6μl Primer F, 0.6μl Primer R, 6.8μl H2O and 2μl template, were run on Applied Biosystems StepOne Plus instrument in triplicate using the following cycle conditions; 95°C for 3 min., 95°C 20 sec., 55°C for 20 sec., 72°C for 30 sec., return to step two 45 times. For standards, full-length bacterial 16S rRNA gene was cloned into a pCR4-TOPO vector, with Kanamycin-Ampicillin resistance. The total plasmid fragment size is expected to be 5556 bp. A bacterial standard was prepared via. 10-fold serial dilutions, and the copies of 16S was determined by the following: Copy# = (DNA wt. x 6.02E23)/(Fragment Size x 660 x 1E9). Linear regression was used to determine copy numbers of samples, based on CT of standards. Reaction specificity was assessed using a melt curve from 55°C to 95°C, held at 0.5°C increment for 1s. For 16s rRNA amplification and sequencing, bacterial 16S rRNA genes were PCR-amplified with dual-barcoded primers targeting the V4 region (515F 5’-GTGCCAGCMGCCGCGGTAA-3’, and 806R 5’-GGACTACHVGGGTWTCTAAT-3’), as per the protocol of Kozich et al. 2013 (Kozich *et al*., 2013). Amplicons were sequenced with an Illumina MiSeq using the 300-bp paired-end kit (v.3). The potential for contamination was addressed by co-sequencing DNA amplified from specimens and from template-free controls (negative control) and extraction kit reagents processed the same way as the specimens. A positive control from samples consisting of cloned SUP05 DNA, was also included. The only modification to this standard protocol was the addition of peptide nucleic acid (PNA) PCR clamps according to the method developed in Lundberg et al. (Lundberg *et al*., 2013). In brief, mPNA, to reduce mitochondria amplification and pPNA to reduce chloroplast amplification, were added into the PCR step during library prep at a concentration of 5uM per PNA. The PCR reaction was then modified with the addition of a PNA annealing step at 78°C for 10s.

### Data analysis

Forward and reverse paired-end reads were filtered and trimmed to 230 and 160 base pairs (bps), respectively using the DADA2 pipeline with default parameters (Callahan *et al*., 2016). Following denoising and merging reads and removing chimeras, DADA2 was used to infer amplicon sequence variants (ASVs), which are analogous to operational taxonomic units (OTUs), and taxonomy was assigned to these ASVs using the DADA2-trained SILVA database. Using DNA extraction and PCR negative controls from 16s sequencing the *decontam* package was implemented using default settings to identify and remove potential contamination from the samples (Davis *et al*., 2018). The assigned ASVs, read count data, and sample metadata were combined in a *phyloseq* object (McMurdie & Holmes, 2013) for downstream analyses. The *phyloseq* package was used to calculate field and greenhouse beta diversity, and a permutational analysis (PERMANOVA) was performed on data rarified to 400 reads (in order to account for the extraordinarily low read count in untreated greenhouse samples) using the *adonis* function in the *vegan* package (Oksanen *et al*., 2022).

All plant growth was analyzed using a linear mixed-effects model in R with the function *lme* from the *nlme* package (Pinheiro *et al*., 2022). Model fit was assessed using ANOVA with the *anova* function to test if the inclusion of additional factors and interactions significantly improved model fit compared to a null model including only the intercept. In each model plant ID was included as a random effect to account for repeat sampling over time.

## Results

### PhylloStart community colonizes the plant phyllosphere

We hypothesized that greenhouse-grown plants would be relatively depauperate in their microbial associations. To test this, we inoculated seedlings with the PhylloStart community by spraying the synthetic microbiome inoculum directly onto the leaves over the course of three, weekly applications. Using an Adonis PERMANOVA we saw no differences in the communities from plants treated with or without Azomite, F = 0.92, p = 0.543, and so for the sake of simplicity these sequences are not included in the figure (Fig. **2**).

**Fig 2.**
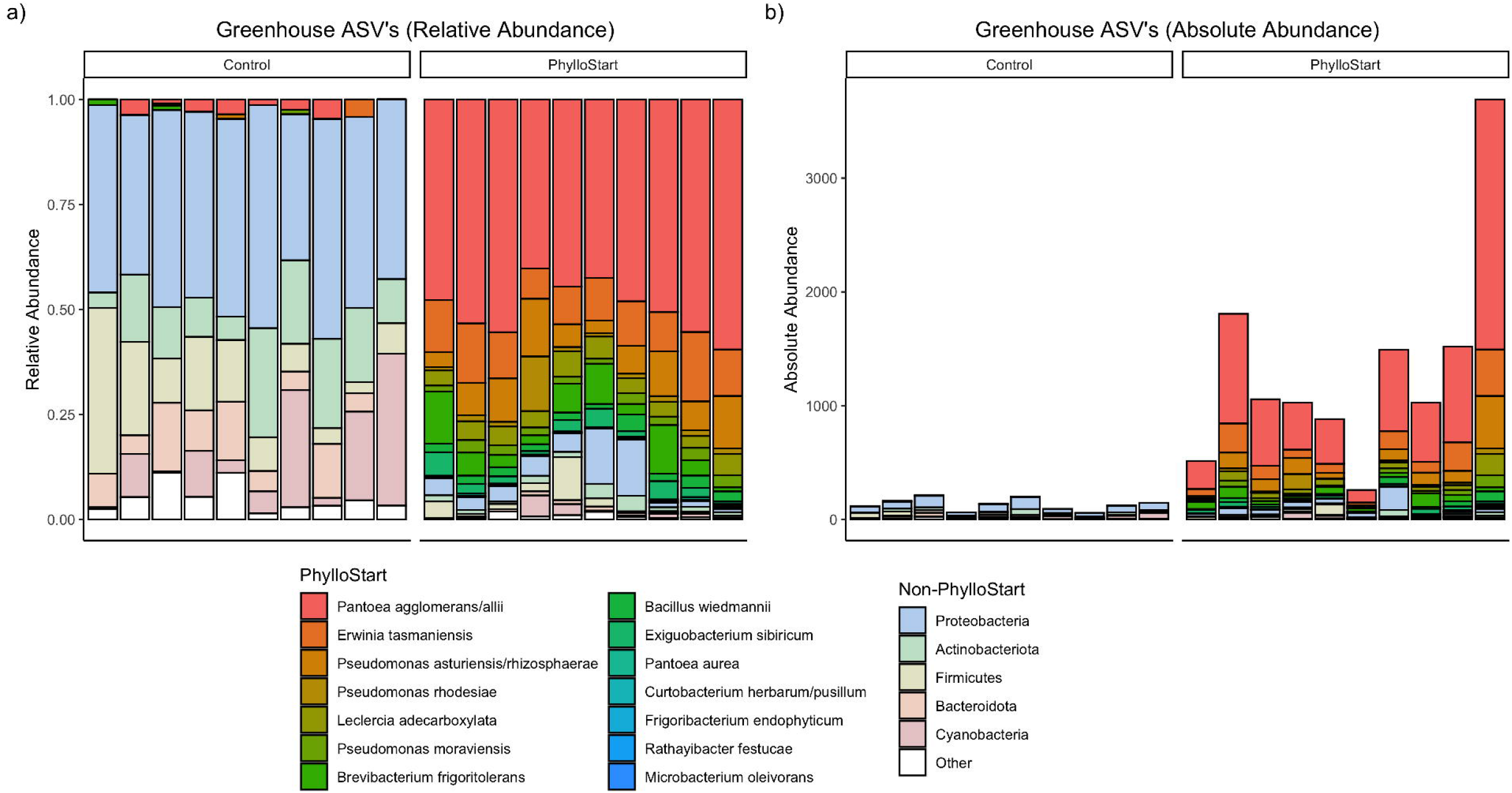
Relative and absolute abundance of ASVs from greenhouse tomato leaves from the first trial. One week after inoculation with the PhylloStart bacteria or a buffer control, only the plants that have been inoculated with PhylloStart have an appreciable number of bacteria residing on their leaves.

We used qPCR to estimate the total number of bacteria on the leaves of PhylloStart treated and control plants, finding that there was a significantly higher abundance of bacteria on the inoculated leaves, t_18_ = −3.97, p = 0.003, with an average of 1327.7 (±950.93 SD) bacterial sequences in the inoculated group, compared to 132 (±52.14 SD) in the controls. Further, in the treated plants, the vast majority of the bacterial sequences were associated with PhylloStart members with an average of 1217.05 (±946.68) PhylloStart-matching sequences per plant. This indicates both that there is robust representation of the PhylloStart on the plant leaves, and that there is minimal development of leaf associated bacteria from the greenhouse environment.

### PhylloStart and micronutrient type interact to increase flowering and fruit production

To determine the role of phyllosphere bacteria, and their interaction with micronutrient supplementation, on greenhouse-grown tomato plants we collected data on a variety of plant characteristics throughout their development. This included plant height, width, flowers, fruit on the plant, and the total weight of fruit harvested. We analyzed the data using a linear mixed effects model, selecting only terms identified by an ANOVA to significantly improve the model’s fit. For height and width, only the model including time was selected as significant (p <0.0001 for both), and unsurprisingly there was a significant effect of time on these traits, as both height and width increase as the plant grows, for height, t_299_ = 106.10, p <0.0001, and for width, t_119_ = 39.26, p <0.0001 (Sup Fig. **1**).

When analyzing flowers and on-plant fruit, we found that the full model (including treatment) was significantly better than the null for both (for flowers, p = 0.0303, and for tomatoes counted on plant, p = 0.007; Sup Fig. **1**). For flowers, we again saw, as expected, a significant effect of time, t_294_ = 13.31, p = <0.0001, as well as a significant impact of the granular and ultrafine Azomite + PhylloStart on number of flowers per plant, t_54_ = −2.32, p = 0.024, with a significant interaction term between this treatment and time, t_294_ = 1.88, p = 0.003 (Sup Fig.**1**). For fruit counted on the plants, we found a significant effect of time, t_292_ = 12.34, p <0.0001, and a significant effect of the interaction between both the PhylloStart only treatment and time, t_292_ = 2.21, p = 0.027, and the granular and ultrafine Azomite + PhylloStart treatment and time, t_292_ = 2.361, p = .019 (Sup Fig. **1**). Given these results, we chose to focus only on the production of fruit in our two subsequent experiments.

### PhylloStart inoculation and micronutrient supplementation increase tomato production

The first experiment fruit was harvested in bulk, preventing the statistical analysis of the resulting harvest data. However, there was a qualitative increase in the total weight of tomatoes harvested from plants inoculated with the PhylloStart bacteria, from 8003.92 grams total yield in control to 9705.54 grams in the PhylloStart treatment and 9302.92 grams in the granular and ultrafine Azomite treatment compared to 10990.6 grams for the granular, ultrafine and PhylloStart treatment (SupFig. **2**).

After establishing that the primary impacts of the PhylloStart bacteria were in flowers (which transition into fruit) and fruit, we repeated the experiment focusing on the total number of tomatoes produced across bacterial conditions and micronutrient supplement. In this experiment, we found a significant increase in the total number of tomatoes produced by plants that were inoculated with PhylloStart bacteria, t_24_ = 3.81, p = 0.001, with an average of 8.89 (±2.93 SD) tomatoes produced per treated plant per harvest, compared to an average of 7.63 (±4.72 SD) tomatoes in the control group (Fig. **3**).

**Fig 3.**
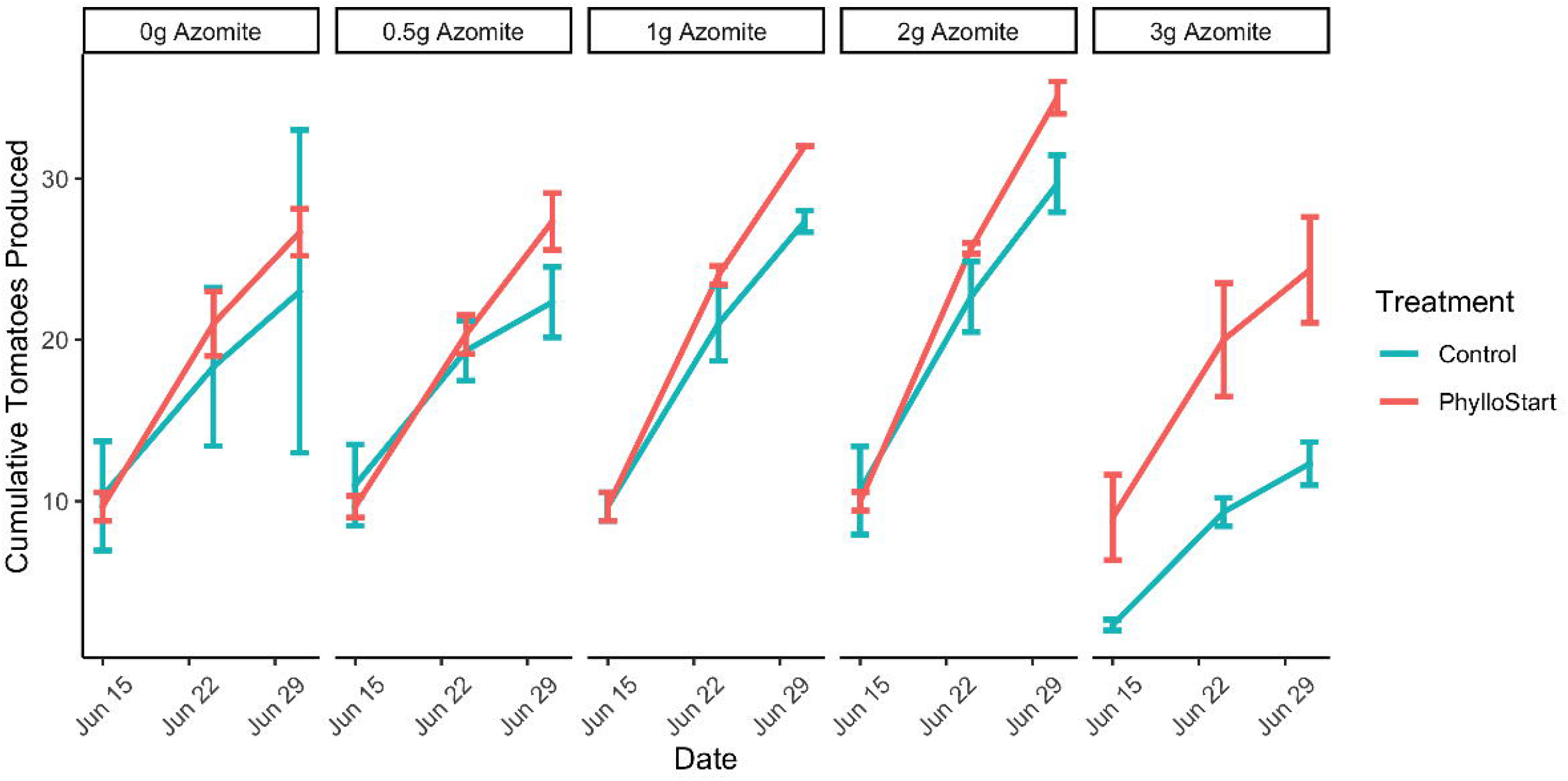
Cumulative number of tomatoes produced across PhylloStart and micronutrient (Azomite) supplemented treatments from the second trial. Both application of the PhylloStart bacteria and Azomite addition lead to a significant increase in the total number of fruit produced. Of note, when adding Azomite in excess (3 grams), total productivity of the control plants was reduced below that of the 0 gram controls. However, when these plants are additionally inoculated with PhylloStart they are rescued to at least the level of the control plants.

Micronutrient supplementation also significantly increased the number of tomatoes produced per harvest, with the 2-gram treatment increasing the average number of tomatoes from 7.63 (±4.72 SD) to 9.89 (±3.41 SD), t(24) = 3.045, p = 0.006. We found no significant interaction between PhylloStart application and Azomite concentration. The highest level of micronutrient supplementation, however led to a significant decrease in yield, with an average 4.11 (±2.32 SD) number of tomatoes produced, t_24_ = −2.18, p = 0.039 (Fig. **3**). As it was clear that the highest concentration of micronutrient supplementation was deleterious to the plant. It is interesting that, while not identified as statistically significant in the model, we observed a trend in which the plants treated with 3 grams of micronutrient supplement and the PhylloStart were less severely affected than those treated with the micronutrient supplement alone (Fig. **3**), indicating that the presence of Phyllosphere bacteria was partially rescuing the plants from this abiotic stress.

In order to rule out that the plants were producing more but smaller tomatoes, we measured both tomato number and weight. Using a linear mixed-effects model, we saw no significant impact of treatment with PhylloStart on the weights of individual tomatoes produced (Sup Fig. **3**). In treated plants, the average tomato weight was 46.05 (±14.99) grams, as compared to an average weight of 46.12 (±11.98) grams in the control group (t_24_ = 1.556, p = 0.133). However, we did see a significant impact of the micronutrient supplement, with 1 and 2 grams significantly increasing the individual tomato weight, t_24_ = 3.83, p = 0.001 and t_24_ = 6.07, p < 0.001 respectively, with average tomato weights of 50.66 (±12.41 SD) grams and 53.46 (±12.39 SD) grams, compared to the control group at a mean of 46.05 (±11.98 SD) grams. Meanwhile, the higher concentration of micronutrient supplementation, at 3 grams per plant, was associated with a significant reduction in the weight of the tomatoes produced, t_24_ = −2.48, p = 0.021, with an average weight of 41.26 (±15.55 SD) grams per tomato. Again, there was no significant interaction between the micronutrient supplementation and PhylloStart bacterial application.

### Phyllosphere amendment increases fruit production in a dose-dependent manner, and these effects persist under disease pressure

To determine if the increased fruit production in PhylloStart inoculated plants was dose-dependent, we repeated the experiment in the fall of 2020, including both the standard inoculum density (OD600=0.02, High) and a lower density (OD600=0.0002, Low). The trends we see in this experiment are consistent with the results from our first and second trials. Notably, these plants were impacted by powdery mildew, a common disease in greenhouse tomato production. The plants began to show signs of infection around week 13, and by week 15 powdery mildew was uniformly present across the surface of most leaves on each plant, regardless of treatment. The plants were randomly dispersed throughout the greenhouse, and regardless of location we did not see a noticeable difference in presence of powdery mildew, so there is no reason to believe that the impacts of disease (beyond the broad impact to the plants as a whole) would bias these results.

Interestingly, despite this disease pressure, the number of tomatoes produced was again significantly increased in the plants inoculated with the PhylloStart bacteria, but only in those treated with the higher inoculation density, t_6_ = 2.70, p = 0.036 (Fig. **4a**). These plants produced an average of 16 (±11.68 SD) tomatoes per plant per harvest, as opposed to 12 (±8.98 SD) tomatoes in the control group. Unlike in the previous experiment, we did not see any significant impact of the micronutrient supplementation on the numbers of tomatoes produced. We also did not see any significant impact of either PhylloStart application (as expected) or micronutrient supplementation (in contrast to our previous experiment) on the average weight of the tomatoes produced (Sup Fig. **3**).

**Fig 4.**
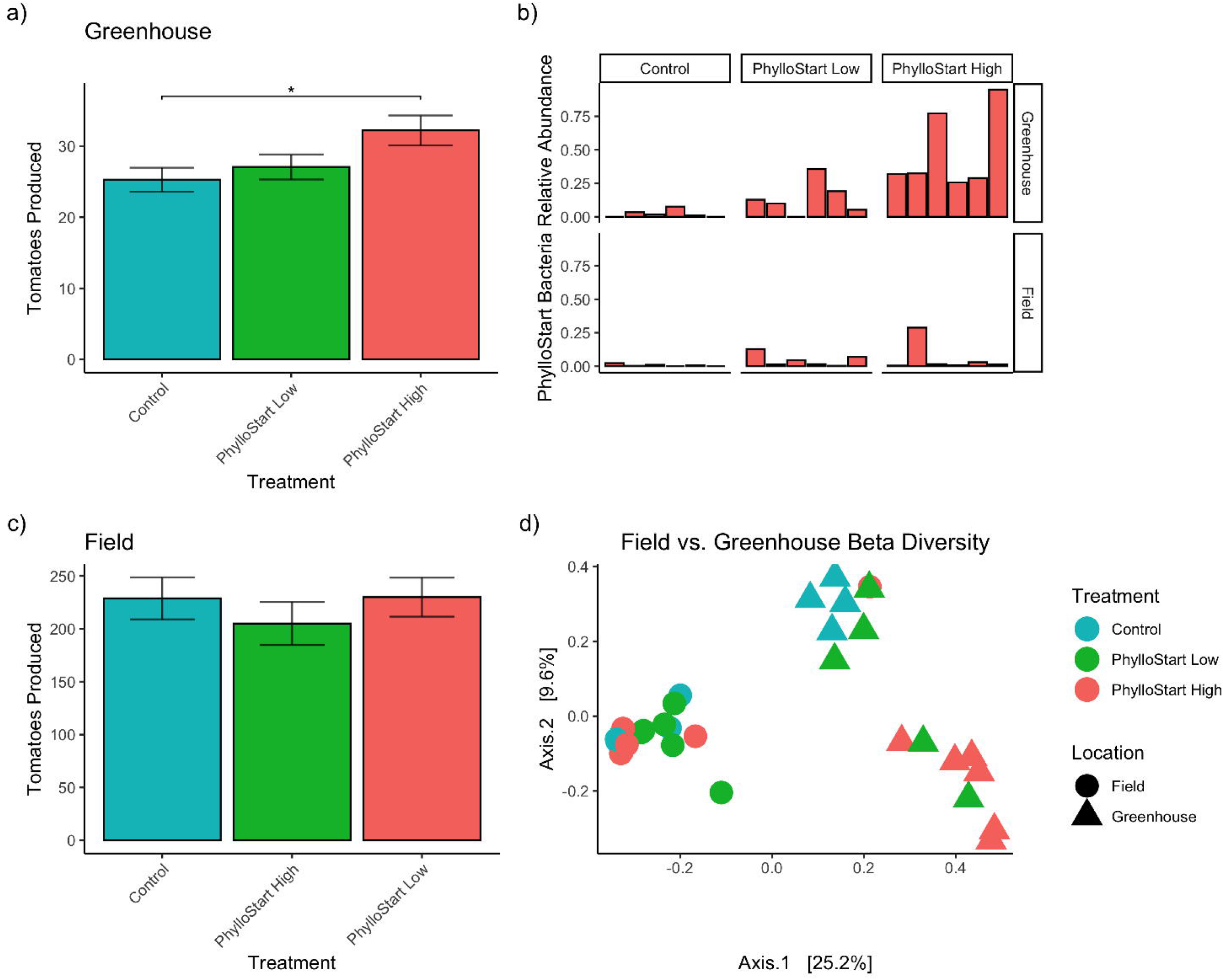
In the third trial greenhouse grown plants show a significant dose dependent effect of PhylloStart, where only the high inoculation density is associated with a significant increase in fruit production. Looking at the relative abundance of PhylloStart bacteria on treated and control plants, one month after inoculation, there are relatively few PhylloStart associated ASVs detected across any of the treatments in the field, while at that time in the greenhouse there are still a marginal number associated with the PhylloStart low density inoculation, and a large portion with the PhylloStart high density inoculation retained on the plants. In contrast to the greenhouse data, there are no significant differences between any of the PhylloStart treatments in the Field grown plants. Finally, looking at Beta Diversity (Bray Dissimilarity), communities group both by treatment and location, with the field communities grouping together, and the greenhouse control and PhylloStart low communities separated from the greenhouse treated PhylloStart high communities.

### Phyllosphere amendment limits subsequent colonization of a bacterial pathogen

Previous work in tomato plants has observed that the native microbiota of the phyllosphere is protective against colonization of the foliar pathogen *Pseudomonas syringae* pv tomato, especially under low resource conditions (Berg & Koskella, 2018). We asked whether the reduced community described in this study would be sufficient to replicate this effect. We applied PhylloStart bacteria to three-week-old plants under either nutrient-limited conditions (grown in autoclaved calcined clay medium) or high nutrient conditions (supplemented with an organic phosphorus fertilizer, Seadbird Guano). One week after PhylloStart bacterial inoculation, leaves were infected with *P. syringae* via blunt-end syringe inoculation. Analysis with ANOVA indicated a significant effect of PhylloStart on pathogen abundance. Post-hoc analysis with a Tukey HSD showed that under nutrient limitation, pathogen load was significantly lower on plants inoculated with PhylloStart bacteria than on plants inoculated with a sterile buffer control, indicating a protective effect, t = 2.67, p = 0.037, however, and as previously observed (Berg & Koskella, 2018), this effect disappeared among plants treated with the phosphorus fertilizer, t = 0.07, p = 0.948 (Fig. **5**).

**Fig 5.**
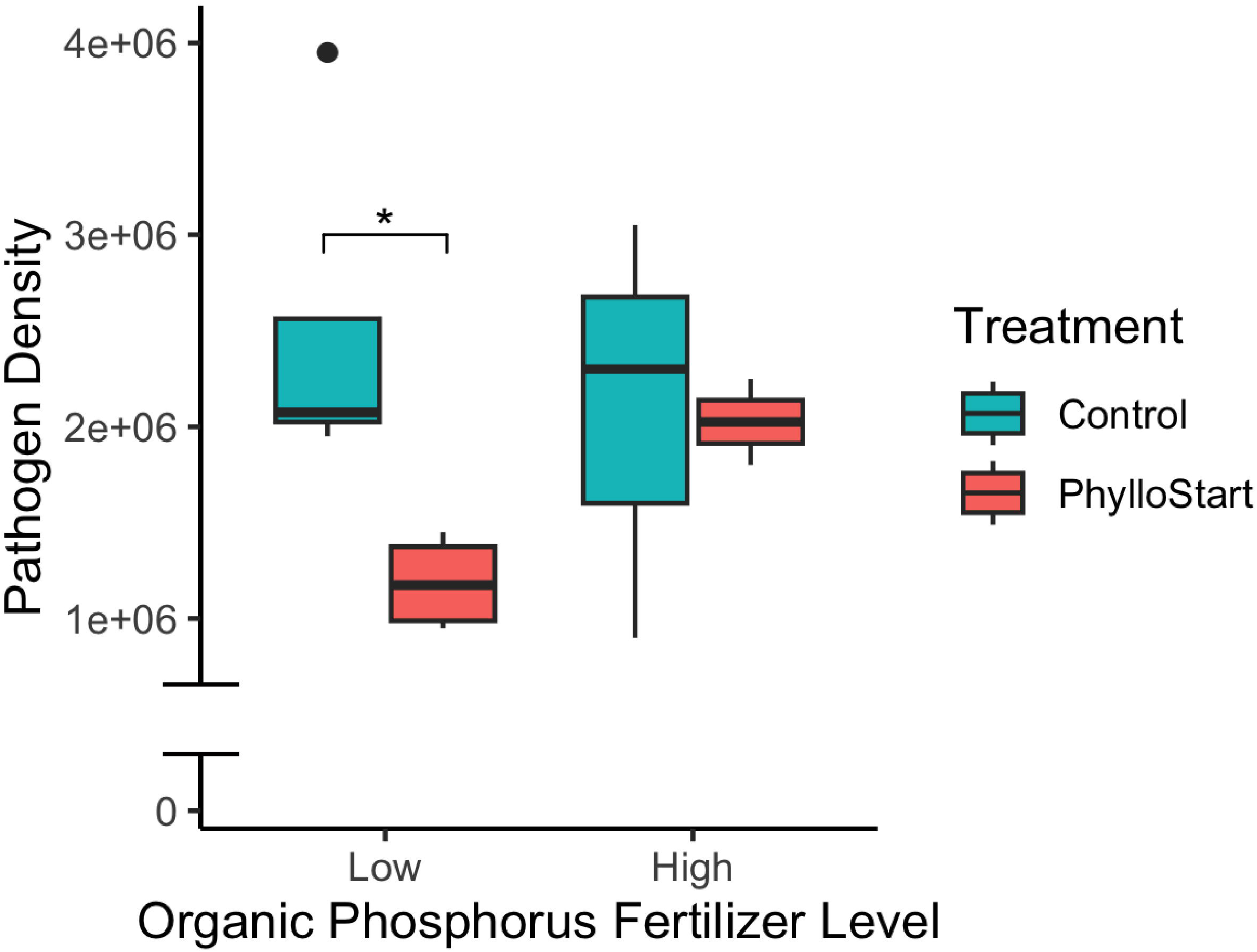
Plants inoculated with PhylloStart bacteria are protected against the establishment of the foliar pathogen P. syringae under low nutrient levels. However, the addition of an organic phosphorus fertilizer leads to a decrease in protection, with both the PhylloStart treated and control plants showing similar levels of pathogen development. Note that, for illustrative purposes, the Y axis is condensed to specific window of density observed on plants and thus does not begin at zero.

### Greenhouse plants maintain PhylloStart bacteria over time, while field plants did not under the conditions tested

In order to determine if the effects of PhylloStart bacteria on plant reproductive success would be seen in an environment with greater dispersal of phyllosphere bacteria and/or whether early inoculation of plants changed subsequent microbiome assembly in the field, we included a field component in the third trial experiment. We transferred both PhylloStart-inoculated and control plants at the end of treatment into the field for the remainder of their development. These plants were sampled concurrently with the plants from the same cohort that remained in the greenhouse (three weeks after their last inoculation), and their phyllosphere communities were sequenced. Using an ANOVA we found that there is a significant effect of inoculation density on the relative abundance of the PhylloStart-associated bacteria in the greenhouse (p = 0.002), which, interrogated with a Tukey HSD, indicated that there was a significantly higher relative abundance in the Phyllostart high treatment compared to the Phyllostart low treatment (p = 0.015) and the control (p = 0.002). Meanwhile, there was no significant effect of PhylloStart treatment on the relative abundance of the PhylloStart-associated bacteria in the field (p = 0.432), (Fig. **4b**).

Furthermore, when looking at a PCOA of community similarity using Bray-Curtis distance metrics (Fig. **4d**) we see that the PhylloStart-treated greenhouse plants clearly separate out from the control plants, with the plants treated with high concentrations of PhylloStart distinct from the controls, and the plants treated with low concentrations of PhylloStart falling between the controls and the high inoculation. Indeed, when analyzing dissimilarity using an Adonis PERMANOVA, we see that the PhylloStart High treated plants are significantly different in community composition from the controls (p=0.004). In contrast, we see no significant difference in terms of PhylloStart relative abundance in the plants that were transplanted to the field, with each treatment containing relatively few of the PhylloStart-associated bacteria. Further, analysis with the Adonis PERMANOVA reveals that there is no significant difference between any of the bacterial treatments in field plants under the conditions tested.

### Early PhylloStart inoculation did not impact field-grown plants

Given that the field-grown plants were able to establish bacteria beyond what was initially inoculated, we sought to determine whether there would be differences in tomato production over the development of the plant. Unlike in the greenhouse experiment, where we found a significant effect of PhylloStart on the total number of tomatoes produced, we did not observe any significant effect of phyllosphere amendment on yield in the field (Fig. **4c**). An ANOVA to test the appropriateness of including PhylloStart in a general linear mixed-effects model, found that inclusion of this factor did not significantly improve the model compared to the null with only the intercept. Like in the greenhouse, we saw no significant effect of PhylloStart on tomato weight (Sup Fig. **3**). In order to verify these results, we performed another field trial in a subsequent year, finding broadly the same results of no significant effect of PhylloStart on the number of tomatoes harvested. In this second experiment we again did not see any significant improvement of the PhylloStart term when compared to the null in a glmm (Sup Fig. **4**). It remains to be seen whether these amendments can provide benefits under broadacre field trials, including under biotic or abiotic pressures where we have observed particularly pronounced effects of PhylloStart in the greenhouse.

## Discussion

This study provides robust evidence that the phyllosphere-associated microbiome enhances the reproductive success of the host plant. Our initial experiment established that greenhouse-grown plants develop a significantly more abundant microbial community when inoculated with phyllosphere native bacterial taxa, and that the effect of early exposure to these bacteria persists throughout the development of the plant. With two additional studies, we verified that these early microbial associations lead to a significant increase in the total number (but not size) of fruit produced by greenhouse-grown tomato plants and that these effects are resilient to both biotic and abiotic stressors. We found these effects to be predominant in the greenhouse setting, as plants that were transplanted into a field environment did not appear to further benefit from the initial inoculation of PhylloStart bacteria. Field performance relies on a combination of additional variables that are commonly controlled in the greenhouse setting. Further field trials under a broad spectrum of conditions and locations are needed to determine whether bacterial amendments to the phyllosphere can potentially confer benefits to commercial field tomato production.

Our qPCR sequencing indicates minimal development of the phyllosphere community in non-treated greenhouse control plants. This supports previous work that found greenhouse-grown plants develop bacterial communities distinct from those in outdoor environments (Maignien *et al*., 2014). This lack of appreciable bacteria in the absence of amendment allowed us to examine the importance of phyllosphere bacteria to plant fitness by inoculating plants with a microbial community designed to mimic phyllosphere communities of field-grown plants. We find that the presence of PhylloStart bacteria, inoculated during the early development of the plant, is associated with increased flowering and fruit production, and that these plants produce a significantly higher amount of fruit throughout their lifetimes. As expected, given microbial dispersal outside of the greenhouse, we did not see these effects persist when transplanting seedlings into a field setting. In this case, PhylloStart-associated bacteria were not found at significant abundances on these plants after a month in the field, and their initial community structure did not seem to shape the future composition of the phyllosphere communities. This may indicate that priority effects during early microbiome establishment are minimally important in a setting with strong colonization pressure from other sources.

There are various mechanisms by which the phyllosphere bacterial community might provide these essential benefits to its host. These primarily include: 1) through altering the plant hormone signaling, either directly through the production of phytohormones or indirectly through the elicitation of a plant response; 2) by increasing the nutrients available to the plant either through enhanced nutrient fixation or availability; and 3) through reduction of stress, either environmental or due to pathogen pressure (Esİtken *et al*., 2005; Paul & Nair, 2008; Adesemoye *et al*., 2008; Beneduzi *et al*., 2012; Bhattacharyya & Jha, 2012). While our current study does not seek to explain the mechanism underlying observed biostimulant effects, it likely relies on a combination of these. However, that the effects of Azomite fertilization and PhylloStart inoculation acted primarily in an additive fashion throughout our study suggests that altered nutrient acquisition as a result of phyllosphere amendment is not a particularly dominant force. Moreover, that the impact of PhylloStart application on yield seemed to be resilient against both the impacts from over-fertilization and disease across our experiments could indicate that these communities play a particularly important role in buffering host plants against stress.

One potential explanation for this increased reproductive success is linked to the phytohormone auxin (or IAA), which is a major regulator of plant growth, is commonly produced by bacteria inhabiting the phyllosphere (Brandl & Lindow, 1998) and has been linked to increased biomass accumulation in rice and corn (Mwajita *et al*., 2013; Abadi *et al*., 2020). In this context, increased fruit yield could be mediated by the action of auxin, decreasing flower abscission (Sexton & Roberts, 1982; Meir *et al*., 2010), potentially leaving more flowers available to set. In support of this idea, we did observe a significant increase in total flowers on the plant over time in our first greenhouse trial (the only one in which this was explicitly measured) in response to PhylloStart application with Ultrafine Azomite (Sup Fig. **1**). Further, using BLAST to search the genomes of the PhylloStart bacteria, we found that several members (*Bacillus wiedmannii*, *Erwinia tasmaniensis*, *Pantoea agglomerans*, and *Pantoea allii*) have matches for *idpC* (indole-3-pyruvate/phenylpyruvate decarboxylase), a key protein in auxin production (Brandl *et al*., 2001). Future work will have to be done to confirm that these bacteria can produce auxin *In Planta*, and if this may explain some of their plant beneficial effects.

It is also possible that the PhylloStart bacteria alter the plant’s response to environmental cues, allowing the plant to better optimize its growth strategy and invest more resources in reproduction. Recent work has focused on the phenomenon of microbiome-dependent ontogenic timing (MiDOT), by which the presence of certain bacterial species acts as essential cues in the developmental timing of their host organism (Metcalf *et al*., 2019). For example, the composition of the *Boechera stricta* (a relative of Arabidopsis) soil-associated bacterial community has been found to significantly alter the timing and duration of flowering. There is further evidence with genetic approaches, such as with the Arabidopsis gene AtBBX32 (Khanna, *et al*., 2009), when introduced into transgenic commercial crops, is responsible for large-scale shifts in the timing of growth responses to external cues like light, which leads to modulation of the timing of reproductive development, including flowering and yield promotion (Holtan *et al*., 2011; Preuss *et al*., 2012). These studies show that the plant is capable of large phenotypic changes triggered by its altered abilities to respond to the incumbent environment, which could be achieved through gene modification or alternative approaches capable of influencing plant physiological responses via the assortment of phytohormones or signals of microbial origin. Further research should continue to assess the role that host-associated microbes play in developmental timing.

Interestingly, we saw that the effects of PhylloStart were particularly pronounced under stressful conditions. In our second trial, where we tested a range of Azomite amounts applied to the soil, it was apparent, in contrast to the benefits at lower concentrations, that the highest concentration of micronutrient supplement produced undesirable results in terms of plant growth and productivity. However, the plants treated with PhylloStart bacteria in addition to this high dose did not see a severe reduction in fruit weight or number of fruit produced, showing performance that was broadly similar to the plants that were not treated with any micronutrient supplement. This effect was apparent again in our third trial, in which powdery mildew severely impacted the plants. In this case, we did not see an impact of the micronutrient fertilizer on the number of fruits produced; however, we still saw a significant impact of association with PhylloStart bacteria on fruit number. Throughout these trials we saw no evidence of a significant direct interaction between the nutrient status of the plant (through micronutrient supplementation) and the effect of the PhylloStart bacteria; instead, these two treatments worked additively to increase the total fruit yield further. Given these observations, we were curious if the PhylloStart community would show the same nutrient dependent pathogen protection as found in our lab’s previous work using a conventional fertilizer (Berg & Koskella, 2018). Indeed, we found that the addition of this community limited the growth of the pathogen *P. syringae* in nutrient-limited plants, but that these effects were abolished when organic phosphorus fertilizer was added. These results are in line with the stress gradient hypothesis, which posits that inter-species interactions should become more facilitative under adverse conditions (Bertness & Callaway, 1994; David *et al*., 2020), and highlight the important role that phyllosphere bacterial associations play in stress response. Further, we see these responses persist well after the initial exposure to PhylloStart bacteria in their early development. This suggests that there may be a critical window in which the plant is receptive to exposure to phyllosphere-associated microbiota, much like what is posited in the hygiene hypothesis for human-associated microbes.

In summary, we find that the presence of phyllosphere-associated bacteria has important benefits to their plant host when they are grown in a microbially depauperate greenhouse environment, primarily through an increase in reproductive success as measured by total fruit production, with further evidence for an increase in stress tolerance. These results are important for understanding the role of microbial communities in host outcomes and are broadly relevant in an agricultural context where, for example, 32% of domestic and 56% of imported tomatoes in the United States are grown in greenhouses that may not provide adequate colonization of phyllosphere bacteria (Baskins *et al*., 2019). Further, we show that bacterial inoculation provides an additive increase in fruit production when applied with a common supplement containing micronutrients, opening avenues for further optimization of agricultural production by harnessing the biostimulant properties of phyllosphere microbes.

## Acknowledgements

This work was funded by NSF EAGER award # 1838299 and CTRI award #2021 - 292. In addition, sequencing costs of plants receiving Azomite treatment were covered in part by AZOMITE®Mineral Products, Inc. This funding did not influence the factual reporting of findings herein. We thank Steve Lindow for excellent discussions on potential mechanisms throughout the writing of the paper, and Gerard Lazo for reviewing different programs for sequence data analysis and presentation. Further, we would like to thank Kama Chock, Isabella Muscettola, Alina Lee, and Fernando Diaz for help harvesting field plants. BK is a Chan Zuckerberg Biohub Investigator and a fellow at the Wissenschaftskolleg zu Berlin.

## Disclaimer

Mention of trade names or commercial products in this publication is solely for the purpose of providing specific information and does not imply recommendation or endorsement by the U.S. Department of Agriculture. USDA is an equal opportunity provider and employer.

## Author Contribution

E.M., K.M., R.D., J.F., R.K. and B.K. contributed to designing experiments; E.M. and R.K. performed research in the greenhouse, all authors were involved in field trials; R.D. and G.K. performed the growth chamber disease assay. E.M. collected samples for sequencing, analyzed data and wrote the paper. K.M., R.D., G.K., J.F., R.K. and B.K reviewed the manuscript and contributed content. B.K. supervised E.M.

## Data Availability

All 16s sequencing data is available on NCBI under the SRA accession number: PRJNA852883.

