## Supplemental Figures for "Early phyllosphere microbial associations impact plant reproductive success"

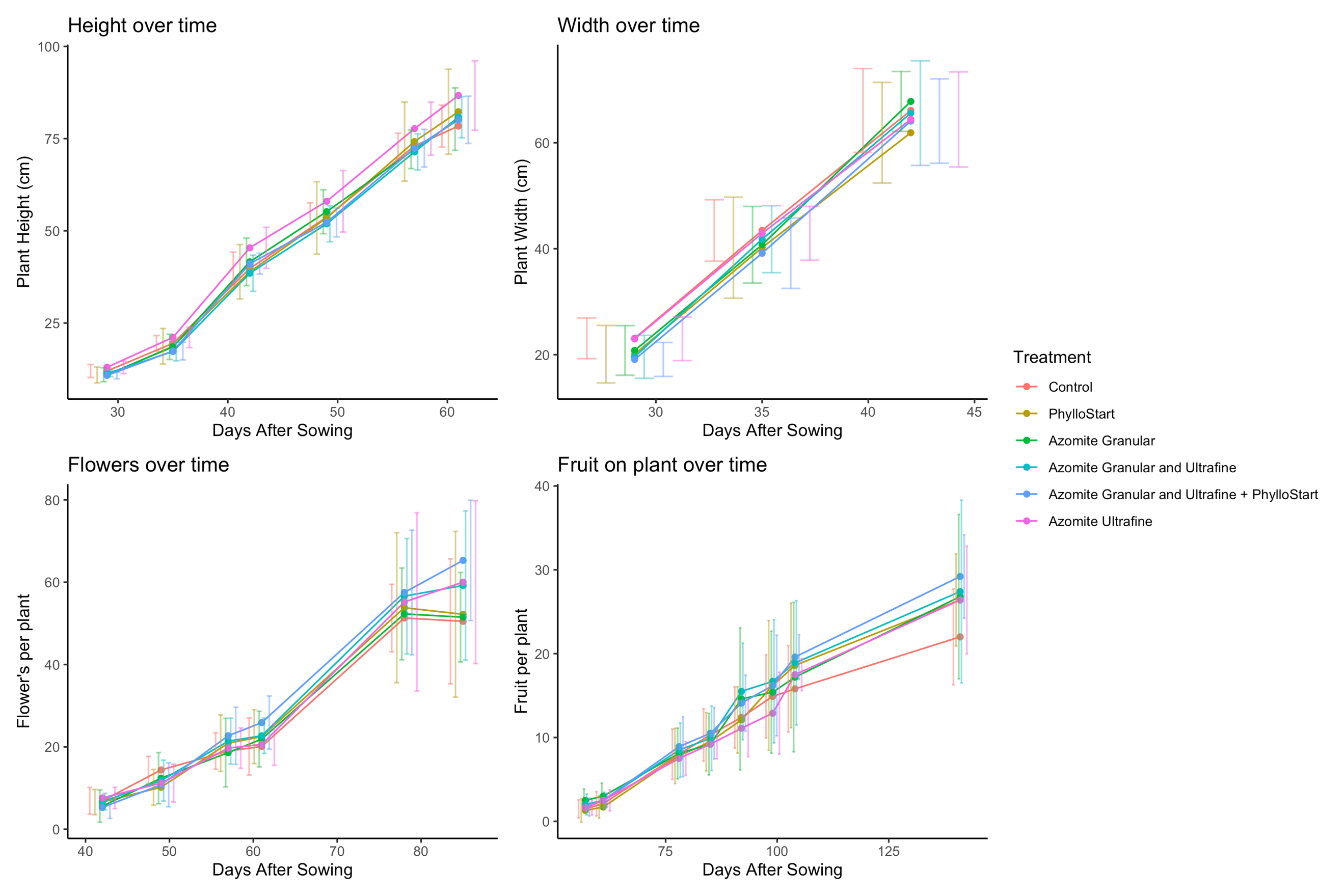

**Sup Fig 1.** Plant metrics measured during the first greenhouse trial. These include plant height, width, flowers, and fruit as measured throughout the growth of the plants. These data were analyzed with a linear mixed effects model selecting only terms that are identified by an ANOVA to lead to a significant improvement in model fit. For height and width only time was identified as a significant factor (p <.0001 and p <.0001 respectively), with height, t(299) = 106.10561, p <.0001, and for width, t(119) = 39.26810, p <.0001, while for both flowers and fruit the full model including PhylloStart and Azomite treatment was found to yield a significantly improved fit (p = .0303 and p = .0074 respectively). Interrogating the full model for flowers there was a significant effect of time, t(294) = 13.314218, p = <.0001, as well as a significant impact of the granular and ultrafine azomite + PhylloStart on flower development, t(54) = -2.317938, p = .0243, with a significant interaction term between this treatment and time, t(294) = 1.881083, p = .0028. Further, for fruit, there was a significant effect of time, t(292) = 12.336256, p <.0001, and significant interactions between both the PhylloStart only treatment and time, t(292) = 2.213625, p = .0276, and the granular and ultrafine azomite + PhylloStart treatment and time, t(292) = 2.361273, p = .0189

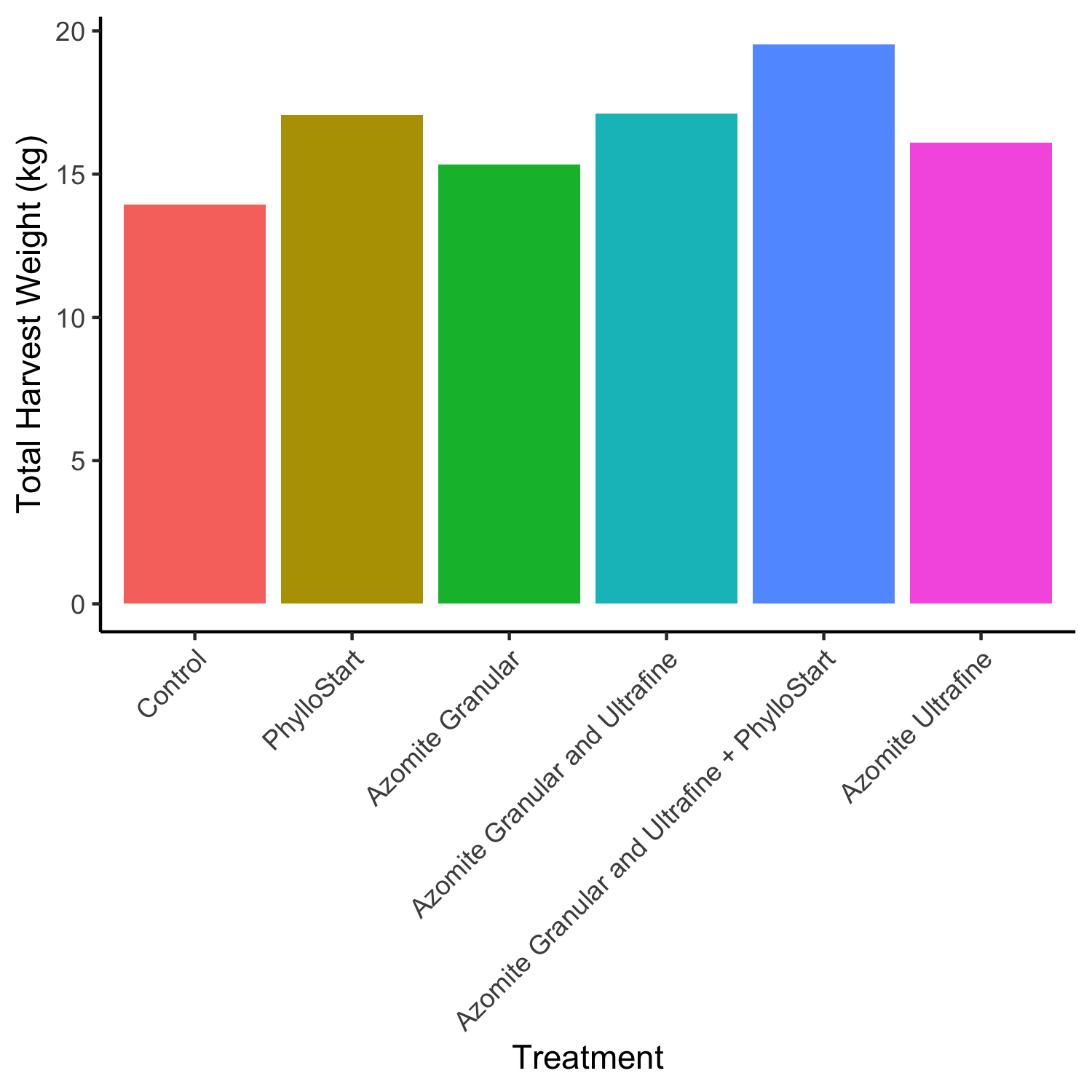

**Sup Fig 2.** Final harvest weights for the initial greenhouse trial. Due to sampling methods, in which each treatment was binned, we cannot statistically analyze the total final harvest. However, we do see a trend where Azomite and PhylloStart treatments yield a larger total harvest, with the combination of Azomite Granular and Ultrafine with PhylloStart showing the greatest increase over the control.

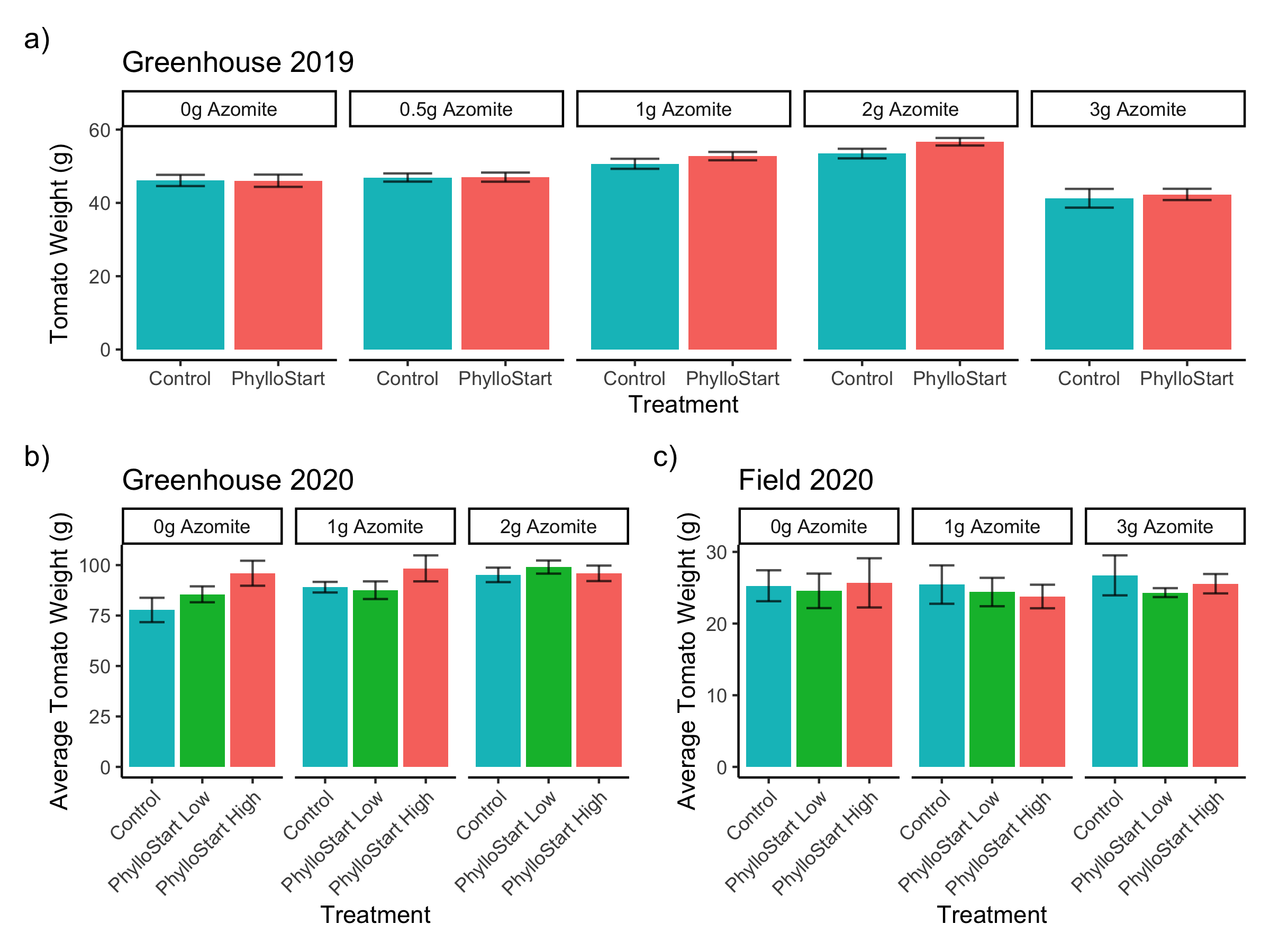

**Sup Fig 3.** Tomato weights as measured throughout the greenhouse and field trials. In the second greenhouse trial, several individual tomatoes from each plant were collected and weighed (a). Analyzing the data with a linear mixed effects model indicated that there was no significant impact of treatment with PhylloStart on the weights of individual tomatoes, t(24) = 1.55582, p = .1328, with treated plants having an average tomato weight was 46.05 (±14.99) grams, as compared to an average weight of 46.12 (±11.98) grams in the control group, there was, however, a significant impact of the micronutrient supplement, with 1 and 2 grams significantly increasing the individual tomato weight, t(24) = 3.83212, p = .0008 and t(24) = 6.07488, p < .001 respectively, with average tomato weights of 50.66 (±12.41 SD) grams and 53.46 (±12.39 SD) grams. In the third greenhouse trial (b), average tomato weight was determined based on the number of total fruit and the total harvest weight per plant. Again, there was no significant effect of PhylloStart on tomato weight, and in contrast to the previous trial there was no significant effect of Azomite treatment. Likewise, there was no significant effect of PhylloStart or Azomite on tomato weight in the field trial (c).

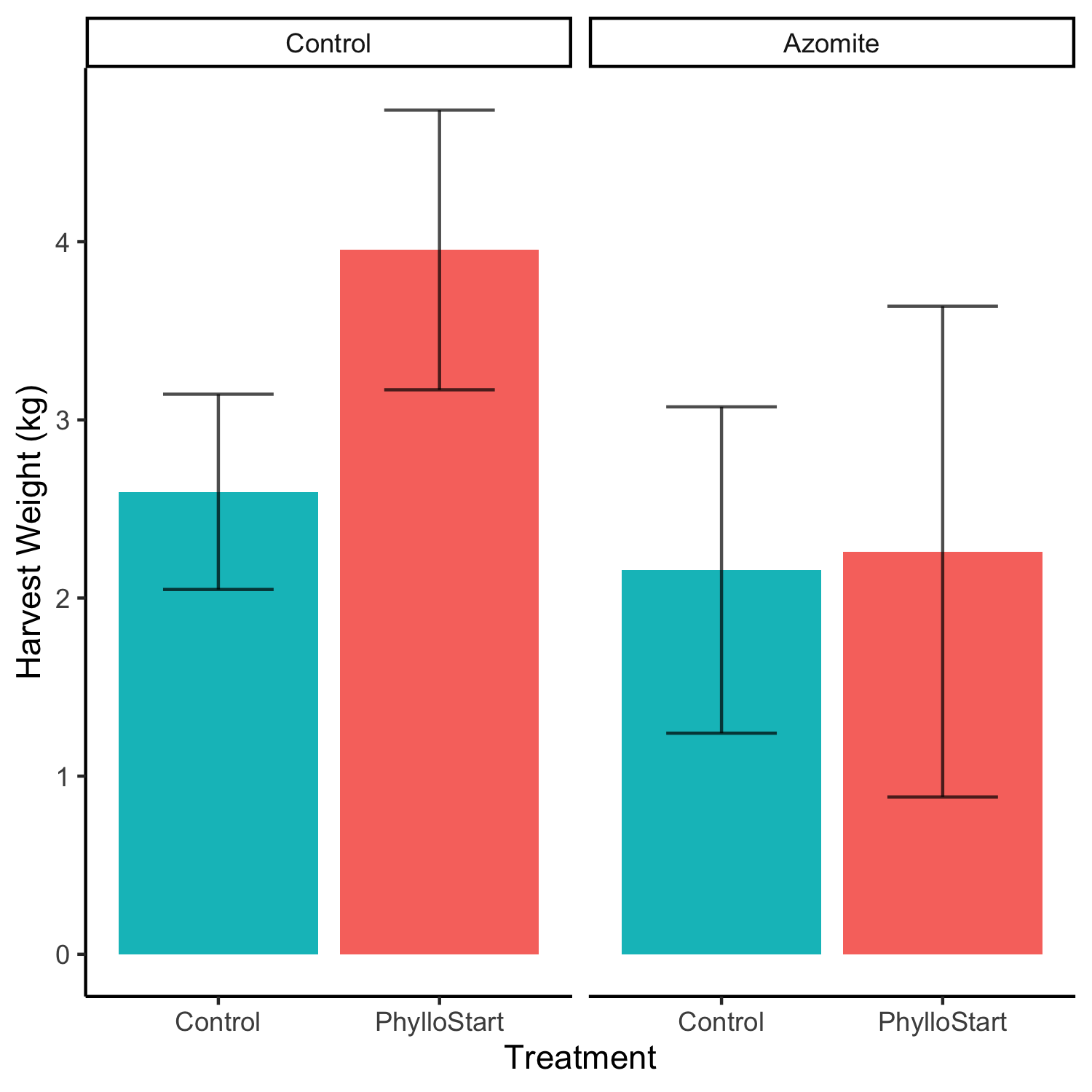

**Sup Fig 4.** In order to confirm that there was no effect of PhylloStart treatment on the number of tomatoes in the field, the experiment was repeated in a subsequent year. Again, there was no significant impact of PhylloStart treatment, or in this case micronutrient supplementation on the total harvest weight at the end of the field season.
